# Contributions of short and long-range white matter tracts in dynamic compensation with aging

**DOI:** 10.1101/2024.02.12.580030

**Authors:** Priyanka Chakraborty, Suman Saha, Gustavo Deco, Arpan Banerjee, Dipanjan Roy

## Abstract

Brain function is shaped by the local and global connections between its dynamical units and biological parameters. With aging, the anatomical connectivity undergoes significant deterioration (e.g., long-range white matter fiber loss), which affects the brain’s overall function. Despite the structural loss, previous research has shown that normative patterns of functions remain intact across the lifespan, defined as the compensatory mechanism of the aging brain. However, the crucial components in guiding the compensatory preservation of the dynamical complexity and the underlying mechanisms remain uncovered. Moreover, it remains largely unknown how the brain readjusts its biological parameters to maintain optimal brain dynamics with age; in this work, we provide a parsimonious mechanism using a whole-brain generative model to uncover the role of sub-communities comprised of short-range and long-range connectivity in driving the dynamic compensation process in the aging brain. We utilize two neuroimaging datasets to demonstrate how short—and long-range white matter tracts affect compensatory mechanisms. We unveil their modulation of intrinsic global scaling parameters, such as global coupling strength and conduction delay, via a personalized large-scale brain model. Our two key findings suggest that (1) the optimal coupling strength and delay play complementary roles in preserving the brain’s optimal working state. (2) Short-range tracts predominantly amplify global coupling strength with age, potentially representing an epiphenomenon of the compensatory mechanism. This mechanistically explains the significance of short-range connections in compensating for the major loss of long-range connections during aging. This insight could help identify alternative avenues to address aging-related diseases where long-range connections are significantly deteriorated.

## 1 Introduction

An overarching question in neuroscience concerns how the brain modulates its biological parameters in response to the age-related deterioration of white matter tracts to preserve functions. Particularly, it is unclear how the two sub-graphs, interacting via short-range (local) or long-range (distant) connections in the brain, contribute to compensation to maintain its desired set point [Deco and Kringelbach, 2016, Deco et al., 2017, Naik et al., 2017, Naskar et al., 2021].

Aging is a complex biological process that is associated with degradation of white matter tracts (a reduction in myelin density, axonal degeneration, and decline in tract numbers) [Peters, 2006, Betzel et al., 2014, Lim et al., 2015, Coelho et al., 2021], gray matter volume [Sullivan and Pfefferbaum, 2006], alteration in neurotransmitter levels [Roalf et al., 2020], and changes in large scale brain networks’ coordination leading to cognitive and behavioral impairments [Li et al., 2020]. It is well known that the impact of aging on brain function is nonlinear and diverse, vary person to person, leading to the deterioration of some functions [Grady et al., 2010], while few other functions, such as inductive reasoning, verbal fluency, and executive attention, may even improve with age [Salthouse, 2012, Veríssimo et al., 2022]. A representative bridge among the aging, its effects, compensation, and the useful measures is shown in Fig. 1.

**Figure 1:**
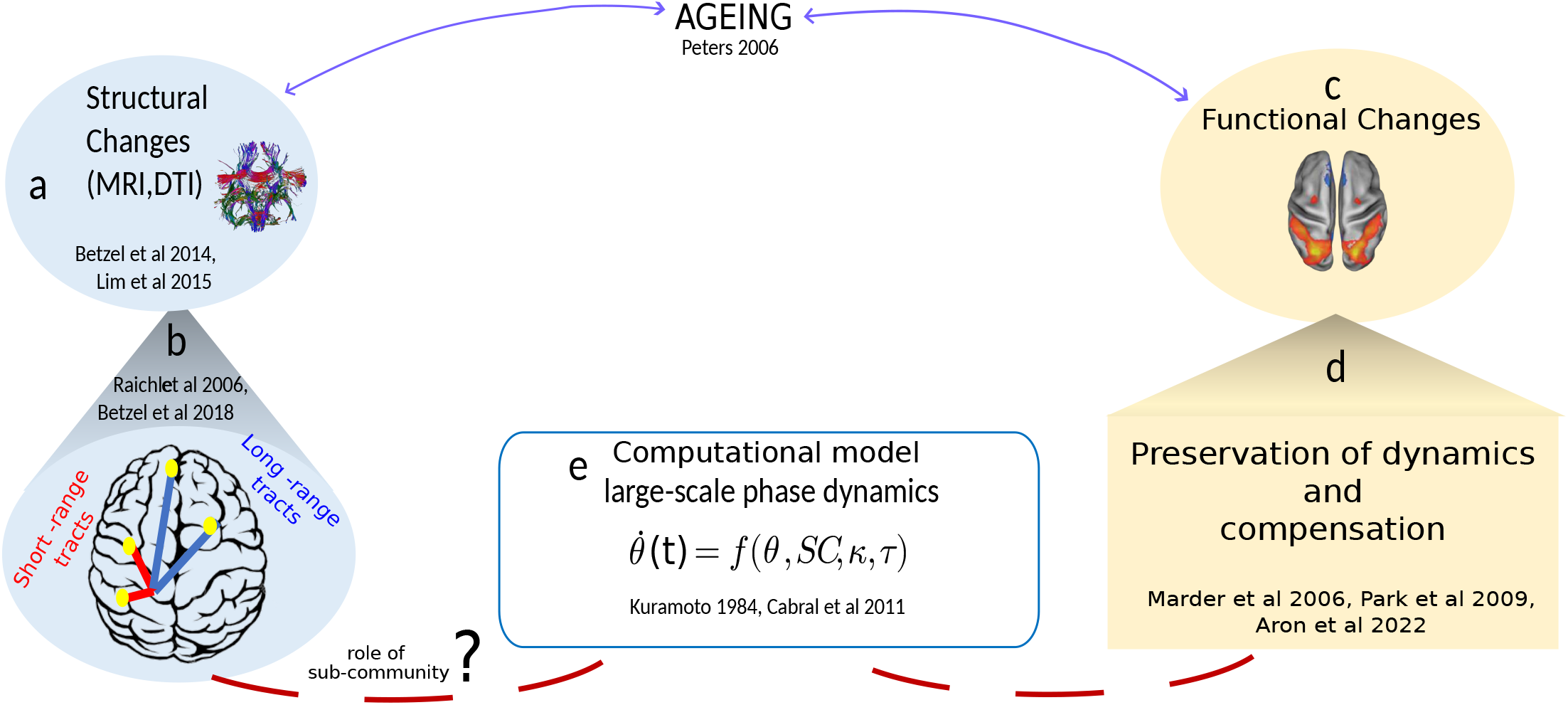
Data analysis and modeling workflow. Figure illustrating significant age-associated alterations in both brain structure and function. (a) indicates a decline in the structural topology of the brain, (b) which includes alterations in both short-range and long-range connections. (c) Studies have demonstrated that the brain undergoes various functional changes, with some functions declining while others remain preserved. (d) Preservation of functions despite structural loss could be interpreted as an age-related compensatory mechanism. (e) Computational models provide a means to study the role of anatomy, specifically white matter tracts such as short-range and long-range sub-communities, in the large-scale dynamics of the brain. In-silico studies have revealed that modulating biological parameters may help preserve dynamic properties of the system, such as synchrony, in the face of structural loss with age. A large-scale phase dynamical model with biophysical parameters, such as global coupling strength and global conduction delay, has been simulated to elucidate the role of the sub-graph in rescaling crucial biophysical parameters coupling strength and conduction delay to preserve dynamical complexity.

The white matter fibers that connect different regions and facilitate efficient communication between neurons undergo significant changes in terms of their length, strength, and myelination as a person ages [Sullivan and Pfefferbaum, 2006, Sullivan et al., 2010, Pannese, 2011], for illustration see Fig 1(a). Short-range connections (or local connections) are the most efficient communication pathways in the brain, as they allow for faster and more accurate transmission of information among closely connected regions [Mišić et al., 2015]. They reduce energy and resource consumption for forming, using, and maintaining brain connections, providing evolutionary benefits across species [Raichle and Mintun, 2006] (Fig. 1(b)). A previous study suggests that short-range white matter connections in the more anterior regions of the left hemisphere may continue to undergo myelination and axonal organization with age [Oyefiade et al., 2018]. Although the brain networks tend to prefer less costly short-range connections, they also have a small number of long (costly) connections, which may provide additional functionality [Avena-Koenigsberger et al., 2017]. The long-range connections (or distant connections) are essential for integrating information across different brain regions by reducing interareal distance [Sporns and Zwi, 2004] and for supporting more complex cognitive processes such as attention, memory, and decision-making [Bassett and Bullmore, 2006, Van Den Heuvel and Sporns, 2011, Ercsey-Ravasz et al., 2013, Bassett and Bullmore, 2017]. Long-range connections have a multifaceted role in brain functions. Betzel et al. demonstrated that long-distance connections play a minor role in reducing topological distance in the brain; instead, they contribute to functional diversity among brain areas, leading to more complex brain dynamics [Betzel and Bassett, 2018]. Additionally, rare long-range exceptions are crucial for improving information processing and the hierarchical organization of the brain [Deco et al., 2021]. Combining short- and long-range connections is essential for ensuring that information is transmitted quickly and efficiently across the brain and supporting a wide range of cognitive functions. A recent study shows that the strength and length of short-range and long-range connections in the brain may decline as we age [Petkoski et al., 2023]. Reduced cortical and subcortical long-range connectivity [Kana et al., 2011] and excess local connectivity is found in ASD [Dajani and Uddin, 2016].

Despite significant structural deterioration with aging, the brain demonstrates a remarkable ability to maintain optimal function, highlighting its adaptability and resilience (Fig. 1(c,d)). Evidence from neuroimaging, cognitive psychology, and neuropathology supports the notion that the aging brain is highly adaptable and resilient [Naik et al., 2017, Uddin, 2021, Aron et al., 2022]. By retaining the ability to adjust and rebound, the aging brain preserves its functional efficiency, compensating for structural alterations [Marder and Goaillard, 2006, Betzel et al., 2014, Naik et al., 2017, Pathak et al., 2022, Petkoski et al., 2023]. This compensatory process, which occurs through synaptic modifications, alternative neural strategies, and adaptive mechanisms such as modulation of biological parameters, is well-explained within theoretical and biophysical frameworks [Park and Reuter-Lorenz, 2009, Pathak et al., 2022,Petkoski et al., 2023] (Fig. 1(e)). Petkoski et al. demonstrated that increased global coupling strength could dynamically compensate for structural loss during aging [Petkoski et al., 2023]. Evidence shows that some level of increased frontal activation with age can be a marker of an adaptive brain that engages in compensatory scaffolding in response to declining neural structures and functions [Park and Reuter-Lorenz, 2009]. Another recent study showed preservation in neural synchrony in aging through enhancing inter-areal coupling [Pathak et al., 2022]. The biophysical mechanism is perfectly plausible from a mechanical viewpoint. However, it still needs to be understood which brain sub-graphs contribute to the compensations and their role in calibrating the global scaling parameters, such as coupling strength and conduction delay.

There is a fundamental knowledge gap regarding how the sub-graphs, connected via short- and long-range tracts, contribute to the compensatory mechanisms in healthy aging while preserving normative functions. This study explores the effects of the two sub-graphs on different biophysical parameters associated with compensation in aging. We try to fill the gap in the role of sub-graphs, parameters, compensation, and aging using the theoretical concepts and their biophysical interpretations. We also investigate the role of short-range and long-range exceptions, which are connections in the human connectome that deviate from the Exponential Distance Rule (EDR) [Ercsey-Ravasz et al., 2013] in the human connectome, in the compensatory mechanisms of the brain. Long-range exceptions, in particular, have been studied for their contribution to enhanced functional information processing [Deco et al., 2021]. However, their impact on the dynamics of the aging brain remains relatively unexplored.

We assume that the changes in anatomical topology in aging, together with the system parameters, control the emergence of collective or coordinated neural dynamics, evaluated under the measure of metastability [Kelso, 2012], as an indicative of optimal working point in healthy aging [Deco and Kringelbach, 2016, Deco et al., 2017, Deco et al., 2019]. We perform in-silico tests using an anatomically constrained phase model (Kuramoto model [Kuramoto, 1984]) with coupling strength (*k*), which globally scales the local interaction strengths, and conduction delay (*τ*), which is associated with the local tract lengths and homogeneously scales the local delay to check the changes in coordinated phase dynamics and observe how the system responds (in-terms of readjustment of parameters and emerged cortical coherence) when the structural topology alters. We justify the choice of this phase dynamical model by focusing only on the phase-dependent measure, i.e., metastability as a key constraining factor in the age-related adaptive mechanisms. Metastability evaluation is based on the phase dynamics obtained from empirical resting-state BOLD fMRI data and model simulations. The metastability captures the dynamical complexity or flexibility of the brain [Kelso, 2012, Deco et al., 2017]. It plays a crucial role in quantifying fluctuations in the instantaneous phase locking of resting-state BOLD signals [Naik et al., 2017], which indicates a healthy brain’s ability to react and respond to an internal or external stimulus promptly. Structural changes associated with aging have been incorporated as input for our model. We systematically manipulate the empirical structural network of individual young subjects and simulate the large-scale generative model with personalized sub-graphs. We validated our findings using two completely independent data sets consisting of 82 subjects from the CamCAN [Shafto et al., 2014, Taylor et al., 2017] and 38 subjects from the Berlin [Schirner et al., 2015] data sets, separated into two groups of young (18-34 years) and elderly (60-84 years) subjects. In neurodegenerative disorders such as multiple sclerosis, Alzheimer’s, and Parkinson’s, both short- and long-range connections in the brain are affected mainly over time [Ouyang et al., 2017, Meijer et al., 2020]. Our results suggest an avenue for using whole-brain models to uncover, in silico, where and how to force the emergence of brain states and may provide an alternative pathway to treat age-related structural degeneracy.

## 2 Materials and Methods

We utilize two datasets for our study: one from the Cambridge Centre for Ageing and Neuroscience (CamCAN) cohort [Shafto et al., 2014, Taylor et al., 2017], and another from Charité University Berlin [Schirner et al., 2015].

### 2.1 CamCAN dataset

#### 2.1.1 Subjects

Our study includes 82 healthy participants (43 females and 39 males) from the Cambridge Centre for Ageing and Neuroscience (CamCAN) cohort. This cohort is available at http://www.mrc-cbu.cam.ac.uk/datasets/camcan/ and has been extensively described in previous studies [Shafto et al., 2014, Taylor et al., 2017]. We divided all participants into two groups: 41 young participants, ranging in age from 18 to 33 (mean age = 25*pm* 4 years, 22 females), and 41 old participants, ranging in age from 60 to 86 (mean age = 74*pm* 6.8 years, 21 females). We referred to these groups as ‘YA’ and ‘OA,’ respectively.

#### 2.1.2 Data acquisition

Resting-state magnetic resonance imaging (MRI), including T1-weighted imaging, diffusion-weighted MRI, and functional MRI, was performed using a 3T Siemens TIM Trio scanner with a 32-channel head-coil at the Medical Research Council (UK) Cognition and Brain Sciences Unit (MRC-CBSU) in Cambridge, UK. High-resolution 3D T1-weighted data were acquired using a magnetization-prepared rapid gradient echo (MPRAGE) sequence with Generalized Autocalibrating Partially Parallel Acquisition (GRAPPA) acceleration factor of 2. Other acquisition parameters were as follows: repetition time (TR) = 2,250 ms, echo time (TE) = 2.99 ms, flip angle = 9°, field of view (FOV) = 256 *×* 240 *×* 192 mm; voxel size = 1 mm^3^ isotropic, inversion time (TI) = 900 ms, and acquisition time (TA) = 4:32 min [Shafto et al., 2014]. Diffusion data were collected using a twice-refocused spin echo sequence with a TR of 9100 ms, TE of 105 ms, FOV of 192 *×* 192 mm, isotropic voxel size of 2 mm^3^, 66 axial slices using 30 directions with b = 1000 s/mm^2^, 30 directions with b = 2000 s/mm^2^, and three b = 0 images with a single average [Shafto et al., 2014]. For resting-state functional MRI, data with an EPI sequence were acquired with a TR of 1970 ms, TE of 30 ms, flip angle of 78°, FOV of 192 *×* 192 mm, voxel size of 3 *×* 3 *×* 4.44 mm^3^, and 32 slices of thickness 3.7 mm, with participants’ eyes closed during acquisition [Shafto et al., 2014].

#### 2.1.3 DTI processing

The diffusion MRI data were processed locally using our pre-processing pipeline, which involved using MRtrix3 ([Tournier et al., 2019], https://github.com/MRtrix3/mrtrix3), FSL (http://www.fmrib.ox.ac.uk/fsl), and ANTS (http://stnava.github.io/ANTs/). The main preprocessing steps included denoising (using the MRtrix command ‘dwidenoise’, [Veraart et al., 2016]), removal of Gibbs ringing artifacts (using the MRtrix command ‘mrdegibbs’, [Kellner et al., 2016]), motion and eddy current corrections (using the MRtrix command ‘dwifslpreproc’, [Andersson et al., 2003]), and bias correction (using the MRtrix command ‘dwibiascorrect’ with ANTs, [Tustison et al., 2010]). Next, a brain mask was calculated in DTI space for each subject using the ‘bet’ command in FSL on the preprocessed image. To find the orientation of the fiber(s) (fiber orientation distribution, FOD) in each voxel, we used multi-shell multi-tissue constrained spherical deconvolution (MSMT-CSD; MRtrix command ‘dwi2response’ and ‘dwi2fod’, [Dhollander et al., 2018]). We then performed a global intensity normalization to make the fiber orientation distributions comparable between subjects. We used Anatomically Constrained Tractography (ACT, MRtrix command ‘5ttgen’, [Smith et al., 2012]) to generate a tissue-segmented image. To use ACT, we preprocessed the T1-weighted image using MRtrix, then co-registered that image to the DWI using MRtrix and FSL. We created a mask of the gray matter/white matter boundary, which was helpful for streamline seeding. Finally, we used probabilistic tractography (MRtrix command ‘tckgen’) to generate 20 million tracts. The tractogram was further filtered (using the SIFT2 approach, MRtrix command ‘tcksift2’) to find a subset of streamlines such that the streamline densities were closer to fiber densities [Smith et al., 2015]. Each step was visually assessed and edited by research personnel.

#### 2.1.4 T1 processing

From, Freesurefer http://surfer.nmr.mgh.harvard.edu [Dale et al., 1999], ‘recon-all’ was employed to reconstruct a two-dimensional cortical surface from a three-dimensional volume acquired from T1-weighted image. The steps involved skull stripping from the anatomical image, estimation of interface between the white matter and grey matte, and generation of white and pial surfaces. The Desikan-Killiany atlas [Desikan et al., 2006] with 68 regions of interest (ROIs) was used for cortical parcellation.

#### 2.1.5 Structural connectivity and fiber tract length

A whole-brain connectome was generated for each subject based on the Desikan–Killiany atlas by computing the fiber density between every pair of regions of interest (ROIs) using the MRtrix command ‘tck2connectome’ with the option ‘scale invnodevol.’ This allowed us to quantify the number of streamlines originating from one ROI and reaching each of the other ROIs. The physical length of the fiber was calculated in millimeters between each pair of ROIs, with a maximum tract length of 250 mm. A tract length (TL) matrix was generated for each subject based on the Desikan–Killiany parcellation.

#### 2.1.6 fMRI preprocessing

Resting-state functional MRI (fMRI) for individual subjects was pre-processed using the CONN toolbox [Whitfield-Gabrieli and Nieto-Castanon, 2012] (https://web.conn-toolbox.org/), which is a Matlab/SPM-based software. The data were pre-processed using the default CONN pipeline with a TR of 1.97 sec. The pre-processing steps included unwrapping using field-map images, realignment to correct for motion, slice timing correction, segmentation, normalization to the MNI template, outlier rejection, and functional smoothing. Spatial smoothing was performed using a Gaussian kernel with a full width of 6.0 mm. Denoising was performed to remove signal changes related to white matter, cerebrospinal fluid, motion, breathing, and cardiac pulsations. Finally, temporal band-pass filtering (0.04-0.9 Hz) and linear detrending were applied. For region of interest (ROI) analysis, the mean regional BOLD time series were estimated in 68 parcellated brain regions of the Desikan–Killiany atlas [Desikan et al., 2006].

#### 2.1.7 Empirical functional connectivity

Aggregated BOLD time series of each region were z-transformed, and then pairwise Pearson correlation coefficients were computed to obtain 68*×*68 resting state functional connectivity (rsFC) matrix for individual subjects.

### 2.2 Berlin dataset

#### 2.2.1 Subjects

We utilized another dataset by Charité University Berlin in [Schirner et al., 2015]. Forty-nine healthy subjects (30 females and 19 males) participated after providing written informed consent to Charité University Berlin, see the Ref. [Schirner et al., 2015]. The age of the subjects ranged from 18 to 80 years, with a mean and standard deviation of 41.55*±*18.44. We divided the subjects into two groups: 24 young participants aged 18 to 33 years (mean=25.7*±*4 years; 13 female) and 14 elderly participants aged 57 to 80 years (mean=70.99*±*9 years; 8 female). The experiments were conducted under the relevant laws and institutional guidelines and approved by the ethics committee of Charité University Berlin [Schirner et al., 2015].

#### 2.2.2 Empirical structural connectivity, tract length, functional connectivity

T1 structural magnetic resonance images (MRI), diffusion-weighted images (DWI), and functional MRI were acquired at the Berlin Center for Advanced Imaging, Charité University Medicine, Berlin, Germany, using a 3T Siemens Trim Trio scanner and a 12-channel Siemens head coil. The voxel size was specified, and the data acquisition tools and pre-processing steps were described in Schirner et al. [Schirner et al., 2015]. Cortical grey matter parcellation was performed using the Desikan-Killiany parcellation method [Desikan et al., 2006], resulting in 34 regions of interest (ROIs) in each hemisphere. Tractography was constrained by seed, target, and stop masks, and the fiber length was represented in millimeters.

The same participants underwent a functional MRI scan during which their eyes-closed, awake, resting-state data were acquired for 22 minutes (repetition time, TR=2 sec). Each region’s pre-processed BOLD time series were z-transformed, and pairwise Pearson correlation coefficients were computed to obtain each subject’s resting-state functional connectivity (rsFC) matrix. All experiments were conducted under the relevant laws and institutional guidelines, and the Charité University Berlin ethics committee approved them.

### 2.3 Defining connectivity based on fiber lengths: SR, MR and LR

There are several definitions available in the existing literature to define short-range (SR), middle-range (MR), and long-range (LR) connections between two ROIs [Wu et al., 2014, Meijer et al., 2020]. Generally, SR connections are intra-regional connections, including intracortical and subcortical connections. MR tracts connect spatially close but distinct regions, including association tracts, commissures, and cortical-subcortical pathways. LR fibers connect spatially distant areas, including inter-hemispheric, cortico-cortical, and cortical-subcortical pathways.

In this study, we defined short, middle, and long-range connections based on fiber tract length setting two thresholds, one at 70 mm and another at 140 mm for CamCAN, and 75mm and 150mm for Berlin, see Table 2. The choice of threshold was based on the approximation of the first (lower threshold for SR) [Meijer et al., 2020] and third (upper threshold for LR) quartiles of the average tract length distribution among young subjects. In Fig. 2(a), we plotted the average fiber tract lengths of the young group. We based our choice of threshold on the approximate first (lower threshold) and third (upper threshold) quartiles of the average tract length distribution. The two blue vertical lines in Fig. 2(a) indicate the lower threshold (e.g., 70mm) and upper threshold (e.g., 140mm), for SR and LR, respectively.

**Table 1:**
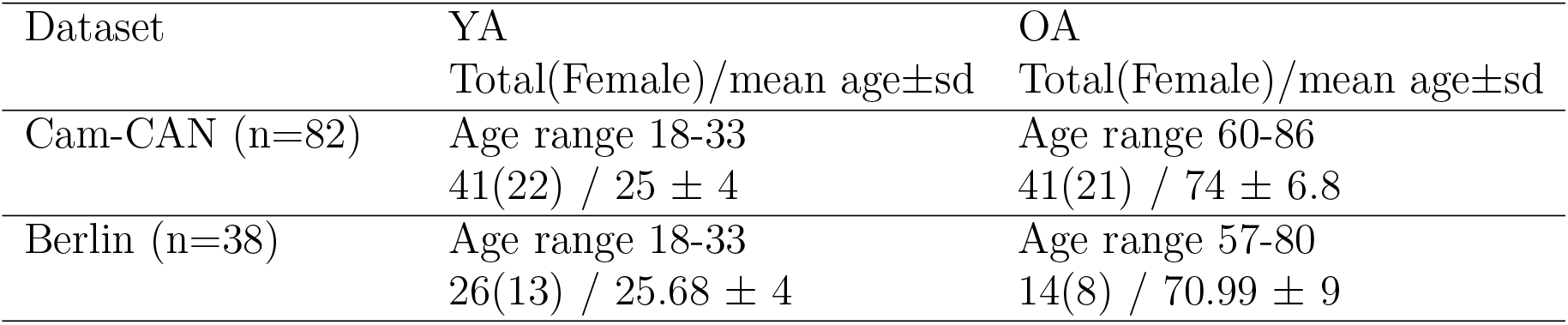
Demographic data of two age groups in two datasets.

**Table 2:** Defining SR, MR, and LR connections for the CamCAN and Berlin datasets.

|  | SR tracts | MR tracts | LR tracts |
| --- | --- | --- | --- |
| CamCAN dataset | tract length $\leq 70$ mm | $70 \text{ mm} < \text{tract length} < 140$ mm | $140 \text{ mm} \leq \text{tract length}$ |
| Berlin dataset | tract length $\leq 75$ mm | $75 \text{ mm} < \text{tract length} < 150$ mm | $150 \text{ mm} \leq \text{tract length}$ |

**Figure 2:**
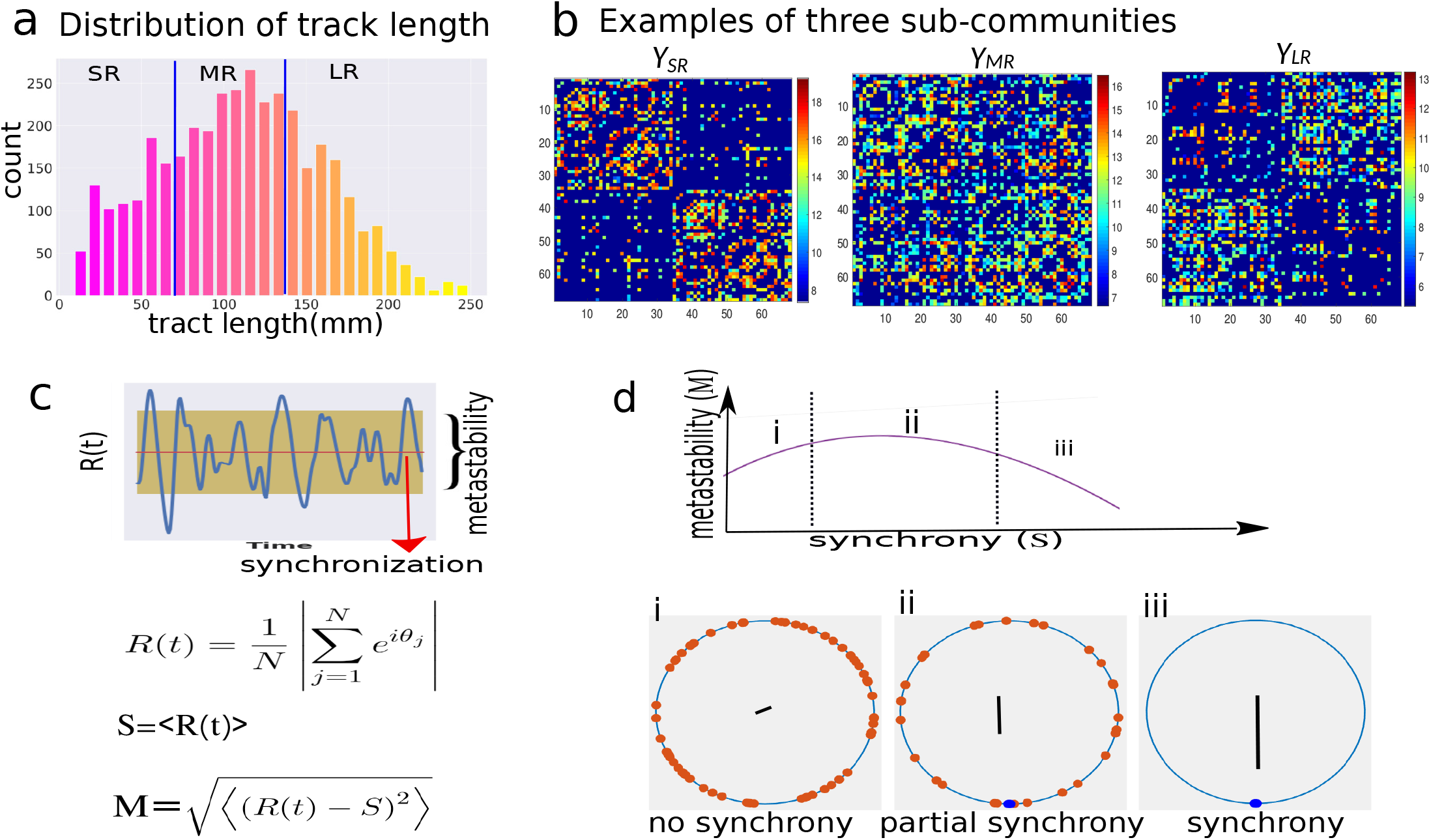
Overview of anatomy and dynamics analysis. (a) The graph shows the average tract length distribution among young subjects of the CamCAN dataset, where the blue vertical lines represent the lower (70mm) and upper (140mm) thresholds. The lengths are categorized into short-range (*SR*), middle-range (*MR*), and long-range (*LR*) based on the thresholds. (b) The intact structural connectivity is separated according to the SR, MR, and LR categories. (c) The Kuramoto order parameter (*R*(*t*)) is plotted against time, where the red line represents the mean of *R*(*t*) that quantifies global synchrony (*S*). The yellow confidence interval indicates fluctuations in *R*(*t*), defined as metastability (*M*) . The analytical expressions of the two dynamical measures are provided below the temporal plot. (d) The relationship between synchrony and metastability is shown in the right panel, which follows a non-linear relationship. The metastability is maximum when the system is partially synchronized. In correspondence to the *M* -*S* characteristic curve, the coherence status is shown for (i) desynchrony, (ii) partial synchrony, and (iii) in-phase synchrony states by the three color-coded phase circle diagrams.

### 2.4 Defining sub-communities comprised of SR, MR and LR connections

The sub-communities or subgraph are defined based on their connection types and combinations, as follows:

i. Sub-community comprised of SR and MR connections without LR connectivity, is denoted as *Y*_*SR*+*MR*_. We keep SR and MR fibers and removed LR connections from the young adult’s structural connectivity (SC), tract length, and functional connectivity (FC).
ii. Sub-community containing only SR tracts, without LR and MR fibers, is labeled as *Y*_*SR*_.
iii. Sub-community, labeled as *Y*_*MR*+*LR*_, connected via MR and LR connections without SR. We retain MR and LR connections while removing all SR connections from the young adults.
iv. *Y*_*LR*_ is the sub-community connected only via LR connections without SR and MR tracts from individual young participants.

An example of three sub-communities generated from a young subject, based on the tract length distribution [Fig. 2(a)], is shown in Fig. 2(b), where the subscripts represent the type of sub-community. Based on the connections types, three exemplary sub-communities are derived from the structural connectivity of young individuals shown in Fig. 2(b). The sub-community connected via SR tracts solely derived from the young SCs, labeled as *Y*_*SR*_ and defined as the SR sub-community of anatomical counts between a given pair of regions. *Y*_*MR*_, is comprised of MR connections only between any pair of regions, shown in Fig. 2(b), middle column. The LR sub-community derived from young SCs is denoted as *Y*_*LR*_ and defined as the sub-community comprised of long-range tracts only, shown in Fig. 2(b), third column.

### 2.5 SR and LR exceptions

Previous research has demonstrated that brain white-matter wiring, based on retrograde tract tracing in non-human primates, can be analytically approximated by the Exponential Distance Rule (EDR) [Ercsey-Ravasz et al., 2013]. In this study, we derived the Exponential Distance Rule for human anatomy using diffusion MRI. Mathematically, the Exponential Distance Rule can be described using an exponential decay function as follows:

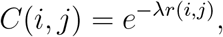

where *C*(*i, j*) is the connectivity weight and *r*(*i, j*) is the distance between *i*-th and *j*-th node. *λ* is the decay. The subject-specific *λ* is obtained from the fitted slope of fiber tract distance versus log weights:

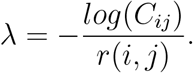

Next, we identified the connections that are exceptions to the EDR rule by focusing on connections that are significantly stronger weights than average, as derived from the EDR. This algorithm first computes the distribution of weight connections at a given distance *r* in the subject-wise connectivity matrix. We then selected only those connection pairs that are three standard deviations above the mean weight of connections at that given distance *r* [Ercsey-Ravasz et al., 2013,Deco et al., 2021]. *These exceptions are categorized based on their tract length range; those within the short-range are termed SR exceptions, while those within the long-range are termed LR exceptions*.

### 2.6 Large-scale generative model: Kuramoto phase dynamics

We selected the Kuramoto model to capture phase transition and global synchrony status with anatomical parameters. The model provides tractable network dynamical measures of whole-brain synchrony and metastability and can capture the key qualitative features of complex brain dynamics. We modeled the dynamical interactions between each *N* (=68) node as a phase oscillator, spatially coupled via structural connectivity [Kuramoto, 1984, Cabral et al., 2011, Breakspear et al., 2010]. The Kuramoto model incorporates two biophysical parameters: conduction delay (*τ*) and global coupling strength (*κ*). Two underlying assumptions are made, i.e., local neural activity is periodic, and the coupling between local neural populations is weak enough to neglect amplitude effects [Daffertshofer and van Wijk, 2011]. The anatomically constrained phase model is computationally less intensive and can simulate microscopic neural dynamics related to underlying structural connectivity [Shanahan, 2010, Cabral et al., 2011, Hellyer et al., 2014, Messé et al., 2014] and emerged collective phase dynamical states.

The dynamics of the *i*^*th*^ phase oscillator (i.e., a brain area) is governed by the Kuramoto phase oscillators as,

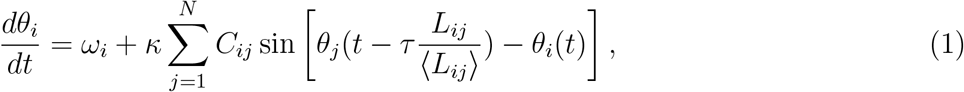

where *i, j* = 1, 2, … *N*. *N* (=68) is the total number of brain regions or the network size. *C*_*ij*_, the normalized connection weights between the *i*^*th*^ and *j*^*th*^ node. *θ*_*i*_ is the phase of *i*^*th*^ oscillator, where *ω*_*i*_ is the intrinsic frequency of oscillation. *κ* is the global coupling coefficient that scales local connection strengths between regions. *τ*_*ij*_ is the conduction delay between node *i* and *j*, defined as *τ*_*ij*_=*L*_*ij*_*/v*, i.e., the ratio of the fiber distance (*L*_*ij*_) and mean conduction speed (*v*) of current through the fibers. For simplicity, we consider the mean conduction delay *τ*, as *(τ)* = *(L) /v*, where *(L*_*i,j*_ ) is the average fiber length. The mean delay can also be interpreted as the global scaling parameters for the local delay between regions. To simplify the model, we assume that the nodes behave as homogeneous neural masses in which all neurons oscillate together. Each node oscillates freely at a fixed frequency of *ω*_*i*_=*ω*=2*π ×* 60 Hz. The model has two free parameters, *κ*, and *τ*. We perform parameter space sweeps to characterize the network dynamics for the entire relevant parameter range. Initial values of the phases are drawn from a normal distribution for a range of *θ*_*i*_(0) ∈ (−*π, π*). We simulate the phase model for a total interval of 80 seconds, where the first 10 seconds of transients are discarded.

Numerical integration of the coupled system was performed using the Euler method with a time step of Δ*t*=0.1 ms. All numerical simulations are performed in MATLAB 2021b (MathWorks). BrainNet Viewer is used for glass brain plots. Python is used to plot other figures.

To correlate the simulated neural activity with the empirically recorded BOLD activity from the subjects, we converted the former to the latter. The fluctuations in firing rate for node *n*, denoted as *r*_*n*_(*t*), fluctuate around a fixed value and are obtained by a periodic function of the local node phase [Cabral et al., 2011]. We chose a simple sine function for these fluctuations, defined as *r*_*i*_(*t*) = *r*_0_ sin(*θ*_*i*_(*t*)), where the amplitude is kept at *r*_0_=1. Next, we used the Balloon-Windkessel hemodynamic model to convert the neural signal to model-based BOLD activity, following methods described in [Cabral et al., 2011].

### 2.7 Dynamical measure: metastability and entropy

We derive metastability for both empirical and computational BOLD signals. The empirical and model-based BOLD signals are converted into a complex phase representation (analytic signal). We compute the analytic signal (*y*_*j*_) from the BOLD series of *j*^*th*^ brain region as,

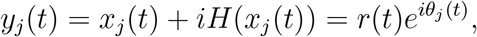

where *H*(*x*_*j*_(*t*)) represents Hilbert transform of the original signal *x*_*j*_(*t*). The instantaneous amplitude and phase of the corresponding BOLD signal are *r*(*t*) and *θ*_*j*_(*t*). A pictorial representation of synchrony and metastability with their definitions is shown in Fig. 2(c).

The Kuramoto order parameter [Kuramoto, 1984], *R*(*t*) is defined as,

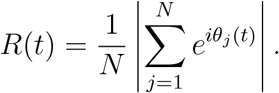

The value of *R*(*t*) quantifies the global coherence status of the network at time *t*, ranging from 0 ≤ *R*(*t*) ≤ 1, where 0 implies no synchrony [Fig. 2(d)i] and 1 indicates a complete or in-phase synchrony state [Fig. 2(d)iii], otherwise partially synchronized [Fig. 2(d)ii]. Mean phase synchrony captures the global coherence status in the network and is defined as,

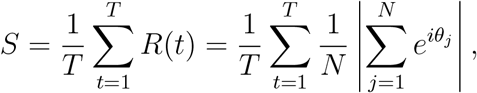

where *N* represents the total number of regions (nodes).

Metastability captures the fluctuations in the synchrony states and is defined as the standard deviation of order parameters over time. The correlation between synchrony and metastability is shown in Fig. 2(d).

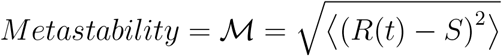

where, ⟨. ⟩ indicates the time average.

In the model simulation, we choose a range of 0.1 to 50 for *κ* and *τ* with an increment interval of 0.6 to capture the metastability map on the *τ* -*κ* parameter plane. The model is simulated to generate the metastability map for each pair of *τ* and *κ* for individual subjects, further utilized to estimate the optimal *τ* and *κ* parameters. Entropy serves as a metric for gauging the intricacy within signals [Bergström and Nevanlinna, 1972, Keshmiri, 2020]. It quantifies the uncertainty or average information content by summing the negative products of probabilities and their logarithms across all potential states. In information theory, it encapsulates the average uncertainty or information content. Mathematically, it’s expressed as

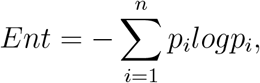

where *p*_*i*_ denotes the probability of each of the *n* states.

### 2.8 Model-based parameter estimation: optimal coupling strength (*κ*) and delay (*τ*)

We utilized a computational framework employing a large-scale brain network model to investigate optimal brain function during a state of maximum metastability at rest [Deco et al., 2017, Surampudi et al., 2019]. The hypothesis of a maximum metastable state is associated with optimal information processing capability and switching behavior of the brain [Deco and Kringelbach, 2016,Deco et al., 2019]. We utilized the maximum metastability condition to compare among the degrees of metastability on the two-parameter plane as a critical constraint to estimate the pa rameters, *κ*^*opt*^ and *τ*^*opt*^. (I) We determined the optimal coupling strength (*κ*) and delay (*τ*) values from the metastability map at the level of individual subject. For each value of the two parameters, we computes *ℳ* (*κ, τ*) to generate a two-parameter space. Next, we determine the optimal parameters utilizing a condition as :

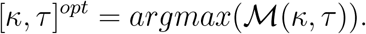

(II) The optimal values of *κ* and *τ* were determined at the individual subject level. The obtained optimal values were subjected to statistical tests to compare young and older age groups.

### 2.9 Group level and subject level comparison: statistical measures

We conducted independent t-tests, using MATLAB built-in function *ttest*2, to compare group differences between young and elderly subjects in the case of CamCAN dataset, and the Mann-Whitney U-test for Berlin data. All statistical analyses were performed using MATLAB, and all reported significance levels are two-sided, with a significance level of *p <* 0.05.

## 3 Results

### 3.1 White matter fiber length and count

We classify the SR, MR, and LR connections using a lower threshold of 70 mm for CamCAN, 75 mm for Berlin, and an upper threshold of 140 mm for CamCAN and 150 mm for Berlin. This enabled us to identify the sub-graphs connected via SR, MR, and LR connections. The average tract length distribution of young and elderly subjects for CamCAN and Berlin cohorts is presented in Figs. 3(a) and 3(d), respectively. On average, there is a reduction of 13% in tract length for CamCAN and 2% for Berlin in the elderly population, which could be due to volume shrinkage [Peters, 2006] or fiber dissociation with age, tabulated in Table 3 for the entire network and other sub-graphs. Figures. 3(b) and 3(e) show scatter plots of subject-wise tract lengths of SR, MR, and LR arranged from left to right panels for CamCAN and Berlin data, respectively. The black lines in Fig.3(b) represent the standard deviations from each group’s mean values of tract lengths. The dashed blue lines connect the mean values of the two groups and exhibit a decaying pattern in tract lengths for elderly subjects. The blue and red dots in the plot correspond to the group’s YA and OA, respectively. The double stars (**) in Fig.3(b) and the single star (*) in Fig.3(e) indicate statistical significance with *p<*10^−6^ and *p<*0.05, respectively. We estimate the percentage of decay in tract lengths for elderly subjects in the CamCAN dataset as 6%, 2%, and 3% and for the Berlin dataset as 3%, 2%, and 0.1% for SR, MR, and LR, respectively (see Table 3).

**Table 3:** Mean (SD) fiber length and fiber strength (count) for the entire network and three sub-graphs, as well as functional network properties for both groups. The percentage of reduction is calculated as the percentage of relative changes in fiber parameters from YA to OA.

|  | YA<br>mean(sd) | OA<br>mean(sd) | Statistics | Percentage of<br>reduction |
| --- | --- | --- | --- | --- |
| **CamCAN dataset** |  |  | (t-test) |  |
| Entire network tract length (mm) | 113 (7.2) | 98 (8) | -7.85 ** | 13% |
| SR tract length (mm) | 43 (1.2) | 40 (1.7) | -8.89 ** | 6% |
| MR tract length (mm) | 107 (1.2) | 105 (2.0) | -6.37 ** | 2% |
| LR tract length (mm) | 171 (3.1) | 166 (4.1) | -4.91 ** | 3% |
| Entire network fiber strength (counts) | $5.1 \times 10^8$ ($1.3 \times 10^8$) | $3 \times 10^8$ ($1.05 \times 10^8$) | -6.54 ** | 35 % |
| SR fiber strength (counts) | $4.3 \times 10^8$ ($1 \times 10^8$) | $3 \times 10^8$ ($0.09 \times 10^8$) | -5.71 ** | 31% |
| MR fiber strength (counts) | $0.54 \times 10^8$ ($0.2 \times 10^8$) | $0.2 \times 10^8$ ($0.1 \times 10^8$) | -10.51 ** | 64% |
| LR fiber strength (counts) | $0.03 \times 10^8$ ($0.01 \times 10^8$) | $0.007 \times 10^8$ ($0.007 \times 10^8$) | -8.19 ** | 78% |
| **Berlin dataset** |  |  | (U test) |  |
| Entire network tract length (mm) | 100 (7.1) | 98 (4.8) | 0.27 *ns* | 2% |
| SR tract length (mm) | 35 (0.5) | 34 (0.5) | 1.21 *ns* | 3% |
| MR tract length (mm) | 94 (0.82) | 93 (0.8) | 1.53 *ns* | 1% |
| LR tract length (mm) | 160 (5.1) | 159 (5.6) | 0.29 *ns* | 0.1% |
| Entire network fiber strength (counts) | $2.9 \times 10^7$ ($0.3 \times 10^7$) | $2.6 \times 10^7$ ($0.2 \times 10^7$) | -3.03 * | 11% |
| SR fiber strength (counts) | $2.2 \times 10^7$ ($0.2 \times 10^7$) | $2.1 \times 10^7$ ($0.2 \times 10^7$) | -1.21 *ns* | 4% |
| MR fiber strength (counts) | $0.72 \times 10^7$ ($0.2 \times 10^7$) | $0.5 \times 10^7$ ($0.1 \times 10^7$) | -3.39 * | 3% |
| LR fiber strength (counts) | $0.02 \times 10^7$ ($0.2 \times 10^7$) | $0.01 \times 10^7$ ($0.1 \times 10^7$) | -2.09 * | 5% |

**Figure 3:**
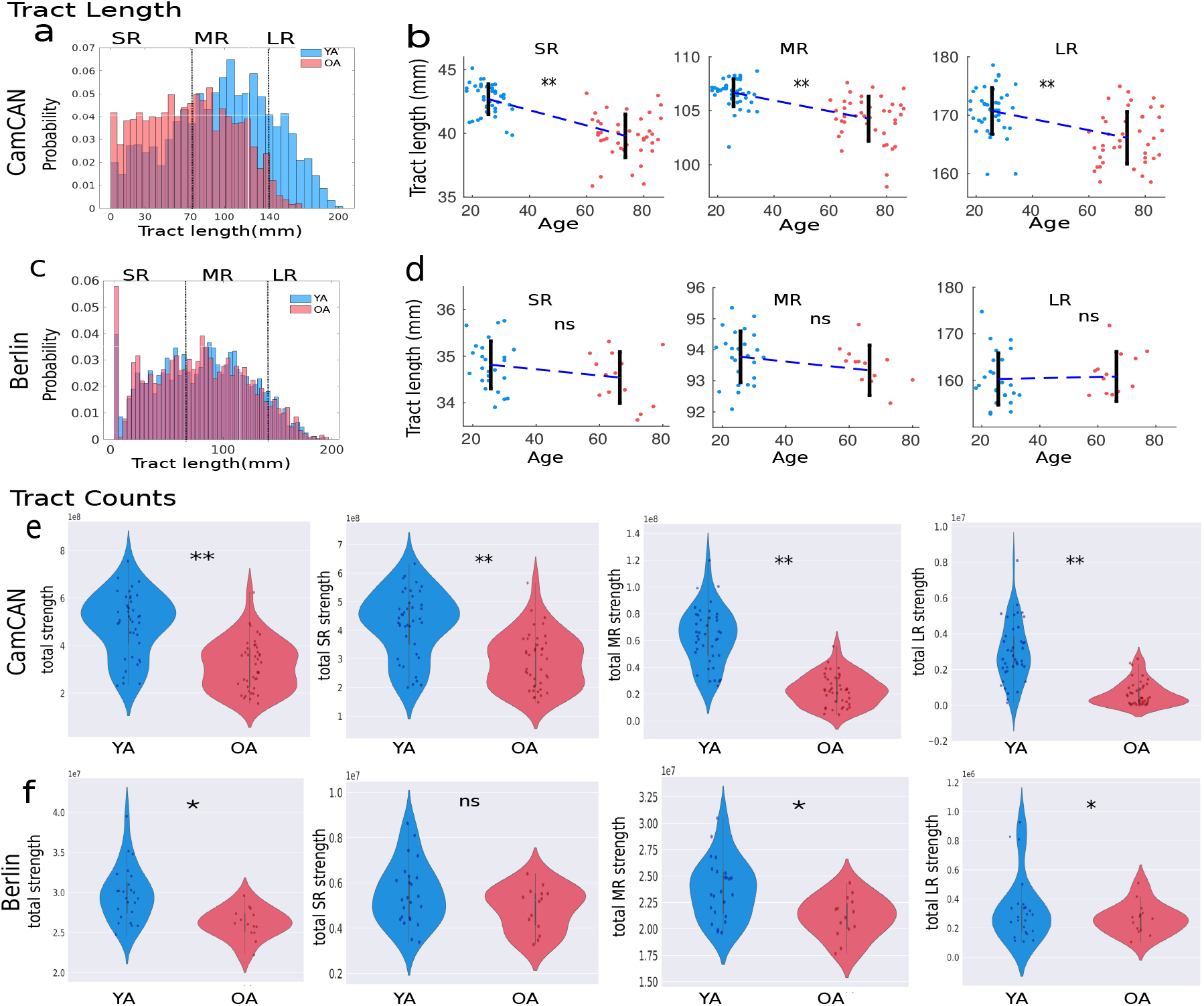
Results from anatomical data analysis shown in (a,b,e) for CamCAN and (c,d,f) Berlin data. (a,c) Distribution of average tract lengths of YA and OA cohorts. (b,d) Distribution of tract lengths (mm) for three sub-communities for YA and OA. (e,f) Distribution of total fiber tract counts of short-range (SR), middle-range (MR), and long-range (LR) connections for the two aging cohorts. Blue and red colors represent YA and OA, respectively.

Next, we examine the reduction in the total number of tracts (counts or strengths). Figures. 3(c) and 3(f) display the violin plots of fiber strengths in the first panel for the CamCAN and Berlin datasets, respectively. Blue and red violins for the YA and OA, respectively. A significant decrease in the total fiber counts is observed in elderly subjects.

Total counts of SR, MR, and LR tracts for young and old subjects are shown in Fig. 3(e) in the CamCAN) and Fig. 3(f) for Berlin datasets. Table 3 tabulates the mean (standard deviation), t-statistic, p-value, percentage of change in fiber length, and fiber strength (count) for the young and old groups. The results show a significant reduction in fiber length and fiber strength (count) in the brains of elderly individuals when compared to their younger counterparts across the entire network and all types of sub-graphs.

SR connections largely dominate the strength of connections in the brain, with approximately 84% of the total connection strength for CamCAN and 78% for Berlin. On average, there is a reduction of approximately 31% in SR tract strength for CamCAN and approximately 4% for Berlin in elderly subjects. In contrast, LR connection strength has minimal overall strength, accounting for only 0.6 % for CamCAN and 0.7 % for Berlin. However, in the elderly group, LR connection strength is found to have deteriorated significantly, with a reduction of 78 % for CamCAN and 5 % for Berlin.

### 3.2 Empirical resting state brain dynamics

We assess the metastability and entropy of large-scale neural dynamics in young and old subjects, measured using 68 regional phase time courses derived from resting-state fMRI BOLD data in both groups. Figures 4(a, b) show the empirical metastability across the two cohorts. Our findings reveal no statistically significant difference in metastability between young and older adults for both the CamCAN and Berlin datasets. We also employ entropy to assess the complexity of brain dynamics. We use a region-based averaging method to compute the overall entropy for a subject. Figures 4(c, d) depict the global entropy across the two cohorts. The entropy values do not exhibit significant differences between the two groups. These results suggest that, at the level of large-scale neural dynamics, the overall stability or variability in the brain’s dynamical repertoire appears invariant in both groups. In other words, despite potential age-related changes in brain structure and function, the fundamental aspect of the brain’s dynamic complexity at a slow time scale remains relatively consistent.

**Figure 4:**
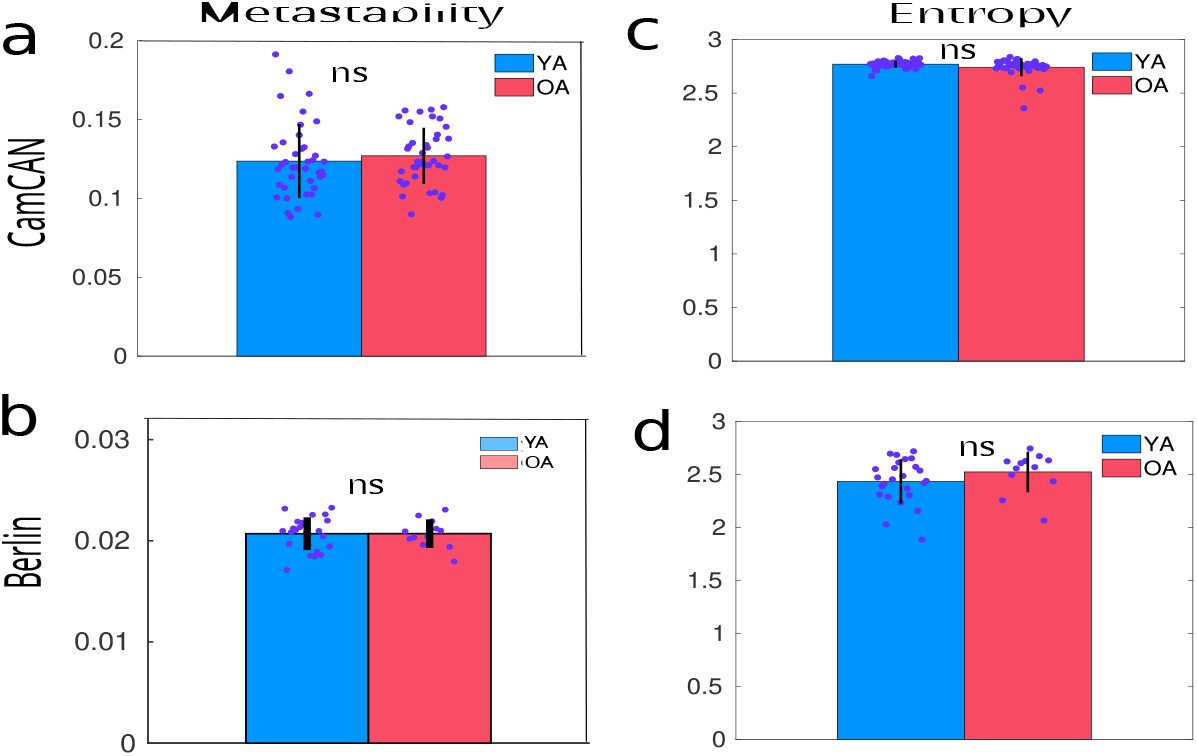
Brain’s dynamical complexity quantified by the metastability (a,b) and entropy (c,d) from resting state fMRI BOLD signals of YA and OA for CamCAN and Berlin cohorts, respectively. Blue and red color represent YA and OA, respectively. *ns* indicates no significant changes between the groups, i.e., *p>*0.05. The consistent level of metastability implies sustained brain dynamics at rest in the two aging cohorts.

### 3.3 SR and LR exceptions on aging

Additionally, we delve deeper to uncover the role of short-range (SR) and long-range (LR) exceptions in the dynamics of the aging brain. First, we apply the EDR rule to the subject-wise connectome, identify the exceptions, and examine how the relative percentage of exceptions changes between young adults (YA) and older adults (OA). Second, we analyze whether these exceptions impact optimal brain dynamics with aging.

Figures 5(a, b) show the relationship between log weights and tract length for each tract, illustrating how the connectivity weight decreases exponentially with increasing tract length. This trend aligns with the Exponential Distance Rule (EDR), which suggests that shorter tracts have higher connectivity weights than longer tracts. Figures 5(c, d) depict the slope distribution of the lines, denoted as *λ*, across subjects in both YA and OA. The slope represents the rate at which the connectivity weight decreases with increasing tract length. There is no significant difference found between the slopes of YA and OA. We identify exceptions by calculating the distribution for a given distance and selecting connection strengths 3 standard deviations above the mean strength distribution. We then calculate the relative percentage of exceptions to the EDR rule for each subject in the CamCAN and Berlin datasets and plot the average value in Figures 5(e, f). This allows us to pinpoint connections that have significantly higher connectivity weights than expected for their length. The exceptions within the short-range (SR) connections range are considered SR exceptions, while those within the long-range (LR) connections are considered LR exceptions. The average percentage of SR exceptions is higher, and the percentage of LR exceptions is lower in older adults (OA) than younger adults (YA) in the CamCAN dataset.

**Figure 5:**
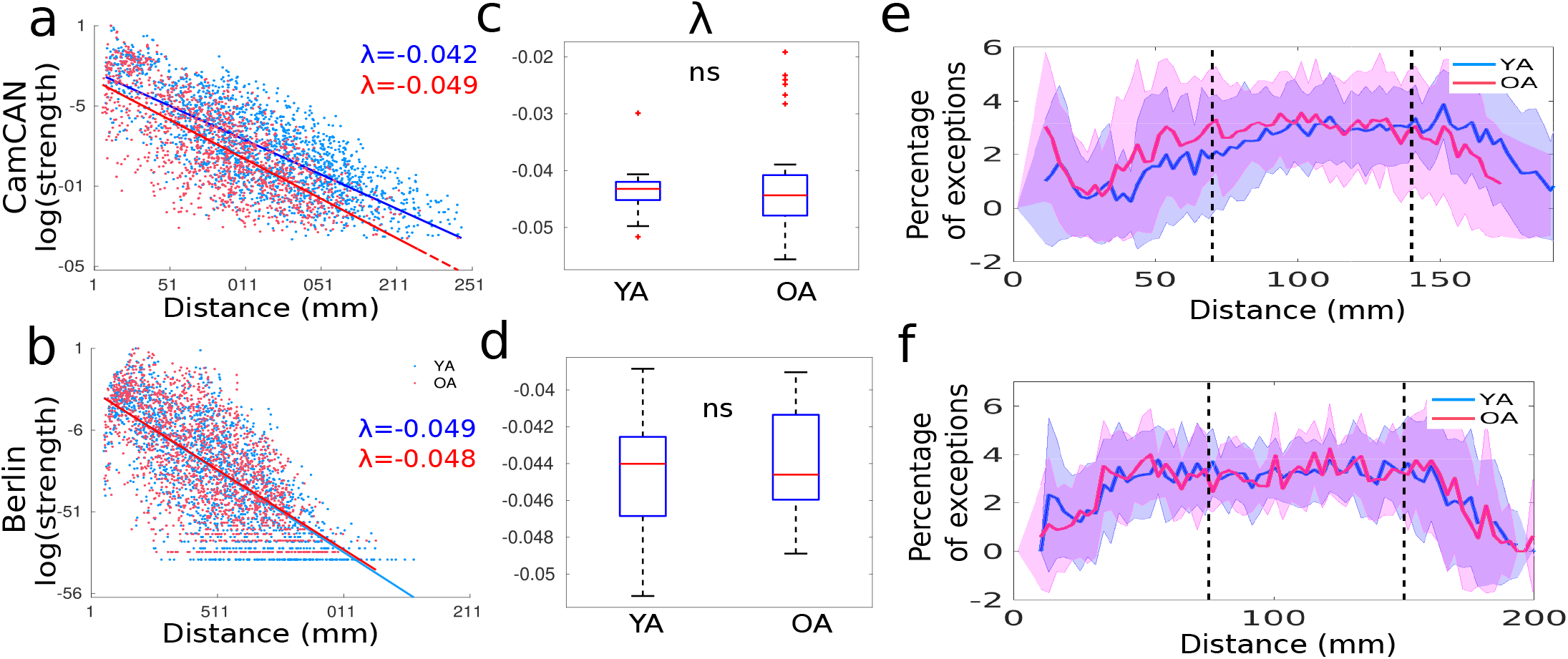
EDR rule and exceptions associated with aging. (a,b) Scatter plot of log weights versus tract length for a subject from YA and OA groups. Each point on the plot represents a specific tract, showing how the connectivity weight decreases exponentially with increasing tract length. (c,d) Boxplot of the slopes, i.e., *λ* over subjects from YA and OA, (e,f) Relative parentage of exceptions in EDR rule for average YA and OA for CamCAN and Berlin datasets. We identify an exception by calculating the distribution for a given distance and selecting connection weights 3 standard deviations above the mean. The percentage of SR exceptions is higher in OA than in YA, whereas older adults have fewer LR exceptions than young adults. Shadow showing the standard deviation of the exceptions over the subjects. ‘ns’: nonsignificant, i.e., *p >* 0.05

### 3.4 Model-based analysis of different the sub-graphs

Our primary analysis focuses on the role of short-range (SR) and long-range (LR) connections in dynamic compensation with aging. However, medium-range (MR) connections may also be crucial for the global dynamical behavior of SR and LR connections. Therefore, investigating the role of SR and LR connections in conjunction with MR connections could lead to more robust and comprehensive findings. Additionally, we explore the role of SR and LR exceptions in large-scale dynamics related to aging. To this end, we create six distinct sub-communities by selectively deleting targeted links from the healthy young’s anatomical connectivity for further analysis. These sub-communities are defined based on their connection types and combinations, as follows:

i. Sub-community comprised of SR and MR connections without LR connectivity is denoted as *Y*_*SR*+*MR*_. We keep SR and MR fibers and removed LR connections from the young adult’s structural connectivity (SC), tract length, and functional connectivity (FC).
ii. Similarly, the sub-community containing only SR tracts, without LR and MR fibers, is labeled as *Y*_*SR*_.
iii. The sub-graph, labeled as *Y*_*MR*+*LR*_, connected via MR and LR connections without SR. We retain MR and LR connections while removing all SR connections from the young adults.
iv. *Y*_*LR*_ is the sub-graph connected only via LR connections without SR and MR tracts from individual young participants.
v. The sub-community containing only SR exceptions and LR exceptions are labeled as 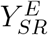 and 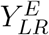, respectively.

Subsequently, we simulate the personalized generative model with the six sub-graphs and intact young and older adults’ structural connectivity. We vary *κ* and *τ* to generate synchrony and metastability maps for individual subjects. We then compare the estimated results of young adults to those of older adults.

Figure 6(a) displays the whole networks of young adults (YA), older adults (OA), and other sub-graphs, including those with both short-range and mid-range connections (*Y*_*SR*+*MR*_), only short-range connections (*Y*_*SR*_), SR exception connections 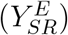, both mid-range and long-range connections (*Y*_*MR*+*LR*_), only long-range connections (*Y*_*LR*_), and long-range exception connections 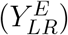. Figure 6(b) illustrates the synchronization (first row) and metastability map (second row) on *τ* -*κ* parameter space for YA, OA, and other six sub-graphs. The region of maximum metastability (depicted in dark red in Fig. 6)) extends in the positive direction of the coupling strength for OA compared to YA. However, the value of maximum metastability (see color bar) remains the same for both groups, indicating modulation of the global coupling strength while achieving maximal metastability.

**Figure 6:**
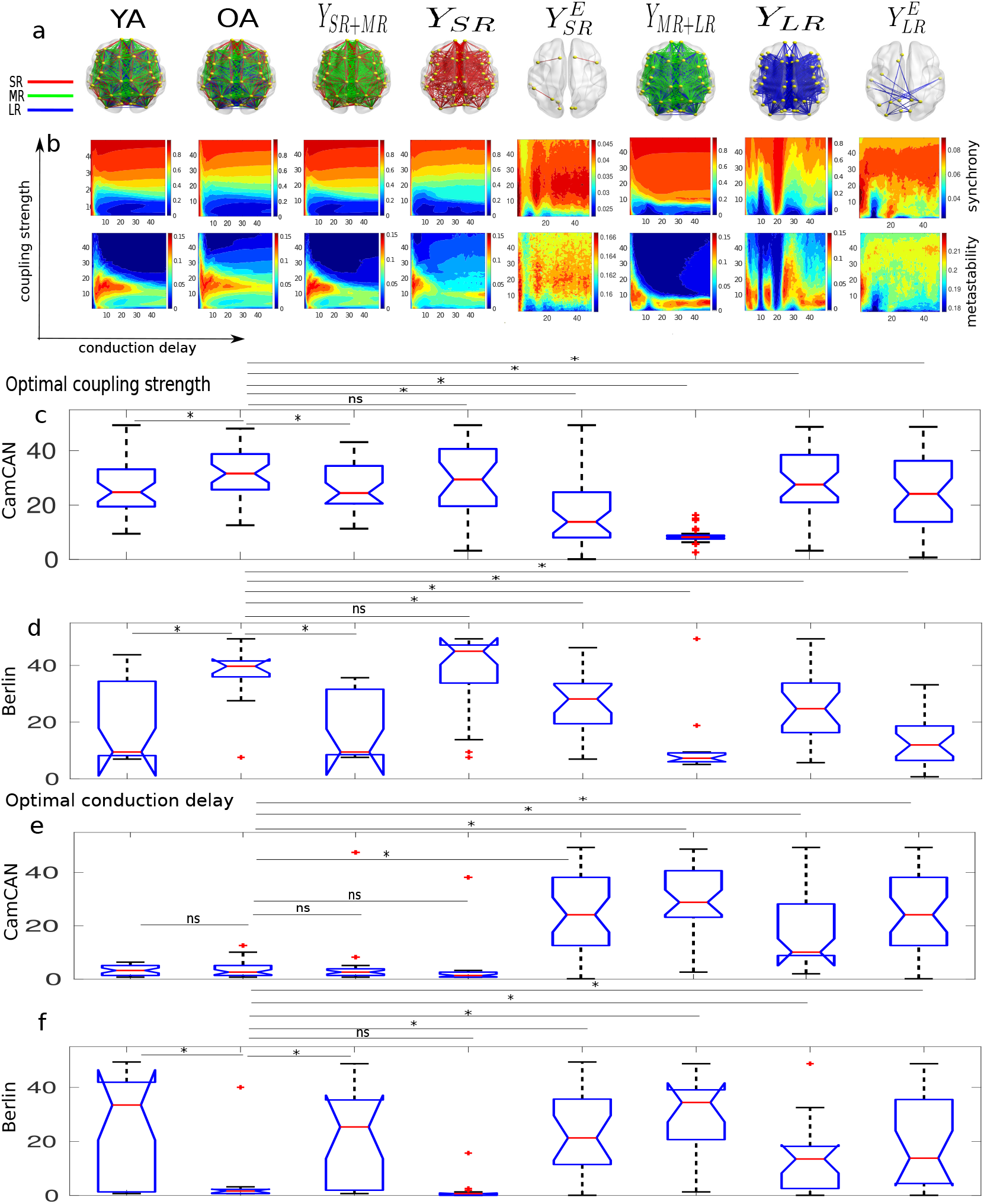
Effects of short-range (SR), and long-range (LR) connections on global brain dynamics. Top panel (a) shows schematic brain views of average tracts for various sub-graphs, including intact networks of young adults (YA), older adults (OA). Plots for the synchronization and metastability maps, optimal coupling strengths, and conduction delays are placed below each sub-group. The second panel (b) displays synchronization and metastability maps for each of the eight cases, generated by taking the average over subjects. Panels (c) and (d) show box plots of model-based optimal coupling strengths estimated from individual subjects from CamCAN and Berlin datasets respectively. We observe significant modulation in coupling strength in OA subjects compared to YA and significant changes between OA and the cases of 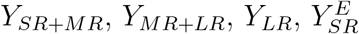, and 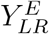 but not with *Y*_*SR*_. Panels (e) and (f) display optimal conduction delay estimates for each case at the individual subject level. Overall, our analysis comparing OA and *Y*_*SR*_ showed no significant changes in the target levels of coupling strength and conduction delay, suggesting that calibration is not necessary when the brain operates with a sub-graph solely connected via short-range connections while operating at maximal metastable state. The two symbols, * and *ns* represent significant (*p<*0.05) and non-significant changes, respectively.

To further assess the similarity between the simulated metastability patterns of OA and the sub-graphs, we average the corresponding maps of each subject and then calculate the pairwise Pearson correlation coefficient between the metastability maps of OA and *Y*_*SR*+*MR*_, OA and *Y*_*SR*_, OA and 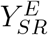, OA and *Y*_*MR*+*LR*_, OA and *Y*_*LR*_, and OA and 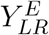. The correlation values are presented in Table 4. The high correlation values observed between the metastability maps of YA and OA indicate that the synthetic metastability patterns remain similar between the two age groups, further supporting our observation on metastability derived from empirical data. To identify which sub-graphs exhibit more similarity in patterns with the OA group, we compared the metastability maps of OA with those of defined networks. When SR connections are present in the network, we observe a strong association between the metastability maps of OA and *Y*_*SR*+*MR*_, as well as *Y*_*SR*_ (Correlation of OA and *Y*_*SR*+*MR*_=0.80 for CamCAN, 0.93 for Berlin; Correlation of OA and*Y*_*SR*_=0.78 for CamCAN, 0.76 for Berlin) . Even correlation between metastability maps of OA and SR exceptions, 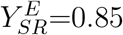 for CamCAN and 0.90 for Berlin. However, when SR connections are absent in the network, specifically for the sub-graphs *Y*_*MR*+*LR*_ and *Y*_*LR*_, the correlation of metastability maps between OA and these sub-graphs decreases (correlation of OA and*Y*_*MR*+*LR*_=0.36 for CamCAN, 0.70 for Berlin; correlation of OA and*Y*_*LR*_=0.46 for CamCAN, 0.56 for Berlin). The correlation between metastability maps of OA and LR exceptions, 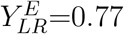 for CamCAN and 0.80 for Berlin. This suggests that SR connections emphasize a consistent metastability pattern similar to the OA, indicating that SR connections might play a significant role in driving the overall metastability of the network for older individuals.

**Table 4:** Correlation between the average metastability maps of OA and the corresponding maps of 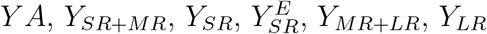, and 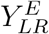 for the CamCAN and Berlin dataset.

| | YA | $Y_{SR+MR}$ | $Y_{SR}$ | $Y_{SR}^E$ | $Y_{MR+LR}$ | $Y_{LR}$ | $Y_{LR}^E$ |
| --- | --- | --- | --- | --- | --- | --- | --- |
| CamCAN dataset | 0.90 | 0.80 | 0.78 | 0.85 | 0.36 | 0.46 | 0.77 |
| Berlin dataset | 0.95 | 0.93 | 0.76 | 0.90 | 0.70 | 0.56 | 0.80 |

To identify the subject-specific biophysical characteristics contributing to maintaining dynamical compensation, global coupling strength (Fig. 6(c,d)), and conduction delay (Fig. 6(e,f)), we select optimal parameters conditioned on maximizing metastability, as described in the Methods section. The mean (standard deviation) values of optimal coupling strength and conduction delay are listed in Table 5. It is noteworthy that when the coupling strength of a subgraph or intact network is low, the conduction delay of that subgraph or network tends to increase, and vice versa. Consider a threshold for the mean optimal *κ* and *τ* set at 25 to clarify this observation. Since the distribution of optimal *κ* and *τ* over subjects ranges from 0 to 50, we choose 25 as the midpoint of the distribution (Fig. 6(c,d,e,f)). We define the mean value of optimal *κ* (and *τ*) over subjects as high if it is greater than 25 and low if it is 25 or below. We can then map high and low values to ‘1’ and ‘0’. By doing so, the mean values from Table 5 can be transformed into a binary format, as shown in Table 6. This table highlights a potential complementary relationship between coupling strength and conduction delay, which could be crucial in maintaining the brain’s optimal working point across different age groups and structural configurations. However, it is noteworthy that the LR exceptions for both datasets do not follow the complementary relationship between optimal strength and delay.

**Table 5:** Mean and standard deviation of optimal coupling strength and conduction delay for different sub-graph types. We perform independent t-tests (for CamCAN) and Mann-Whitney U tests (for Berlin) to compare the optimal parameters between young adults (YA) and older adults (OA), as well as between OA and other sub-graphs. Significant differences are denoted by a star (^∗^) in the superscript of the corresponding group, while non-significant changes are indicated by *ns*.

| | YA<br>mean(sd) | OA<br>mean(sd) | $Y_{SR+MR}$<br>mean(sd) | $Y_{SR}$<br>mean(sd) | $Y_{SR}^E$<br>mean(sd) | $Y_{MR+LR}$<br>mean(sd) | $Y_{LR}$<br>mean(sd) | $Y_{LR}^E$<br>mean(sd) |
| --- | --- | --- | --- | --- | --- | --- | --- | --- |
| CamCAN dataset |  |  |  |  |  |  |  |  |
| optimal $\kappa$ | 26.59 (10.1)* | 32.11 (9.6) | 26.33 (8.5)* | 30.10 (11.4)<sup>ns</sup> | 17.18 (10.1)* | 8.50 (2.6)* | 27.34 (13.4)* | 17.18 (10.7)* |
| optimal $\tau$ | 3.46 (2.0)<sup>ns</sup> | 3.53 (2.8) | 2.69 (1.6) <sup>ns</sup> | 2.72 (6.4)<sup>ns</sup> | 23.10(12.1) | 30.0 (12.1)* | 18.84 (15.6)* | 23.52 (6.5)* |
| Berlin dataset |  |  |  |  |  |  |  |  |
| optimal $\kappa$ | 19.25 (13.1)* | 37.26 (10.1) | 17.80 (11.3)* | 38.31 (13.2)<sup>ns</sup> | 27.05 (9.1)* | 10.51 (10.8)* | 25.84 (11.4)* | 13.22(7.2) * |
| optimal $\tau$ | 24.34 (11.1)* | 4.72(11.1) | 21.08 (17.0)* | 2.41 (10.1)<sup>ns</sup> | 22.31(11.2)* | 29.33 (15.2)* | 13.68 (11.8)* | 19.42(8.2)* |

**Table 6:** Complementary relation between optimal *κ* and optimal *τ* over the groups. ‘1’ indicates that the mean optimal values (*κ, τ*) over subjects are greater than the threshold, 25, and ‘0’ indicates that they are less than equal to 25.

| | YA | OA | $Y_{SR+MR}$ | $Y_{SR}$ | $Y_{SR}^E$ | $Y_{MR+LR}$ | $Y_{LR}$ | $Y_{LR}^E$ |
| --- | --- | --- | --- | --- | --- | --- | --- | --- |
| CamCAN dataset |  |  |  |  |  |  |  |  |
| optimal $\kappa$ | 1 | 1 | 1 | 1 | 0 | 0 | 1 | 0 |
| optimal $\tau$ | 0 | 0 | 0 | 0 | 1 | 1 | 0 | 0 |
| Berlin dataset |  |  |  |  |  |  |  |  |
| optimal $\kappa$ | 0 | 1 | 0 | 1 | 1 | 0 | 1 | 0 |
| optimal $\tau$ | 1 | 0 | 1 | 0 | 0 | 1 | 0 | 0 |

Figures 6(c) and 6(d) for the CamCAN and Berlin cohorts show the boxplots for optimal coupling strength, respectively. We observe a significant increase in the optimal coupling strength (*κ*) for older adults compared to younger groups in both datasets (*F* (77)=2.34, *p<*0.05 for CamCAN; *Z*=2.44, *p<*0.01 for Berlin). Increasing the coupling strength may be an adaptive process associated with the dynamical compensation for age-related long-range white-matter fiber loss.

Further, to investigate which specific sub-graph increases coupling strength in elderly cohorts, we compare the optimal *κ* estimated from the OA group with that of other sub-graph types. The optimal *κ* for 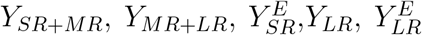 significantly decrease compared to OA (CamCAN: *F* (69)=-2.65, *p<*0.05 for *Y*_*SR*+*MR*_; *F* (65)=-2.2, *p<*0.05 for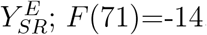, *p<*0.05 for *Y*_*MR*+*LR*_; *F* (70)=-2.26, *p<*0.05 for *Y*_*LR*_; *F* (65)=-2.06, *p<*0.05 for 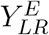; Berlin: *Z*=3.17, *p*=0.001 for *Y*_*SR*+*MR*_; *Z*=2.55, *p*=0.03 for 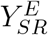; *Z*=3.55, *p*=0.003 for *Y*_*MR*+*LR*_; *Z*=2.27, *p*=0.02 for *Y*_*LR*_; *Z*=2.1, *p <*0.05 for 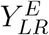). However, we find no significant difference between the optimal *κ* of OA and *Y*_*SR*_ (for both datasets), indicating that the optimal *κ* of sub-graphs containing SR connections is similar to that of optimal coupling strength of older adults. Figure 6(e) for the CamCAN and Fig. 6(f) for Berlin cohorts show the box plots for optimal conduction delay (*τ*) . We observe that the optimal conduction delay for the entire network remains unchanged for YA and OA groups in CamCAN. In the Berlin dataset, the delay value is very high in YA compared to OA (*Z*=2.25, *p <* 0.05). We find no significant difference between OA and *Y*_*MR*+*SR*_ for CamCAN and a substantial difference for Berlin (*Z*=2.62, p*<* 0.5). Also, there is no significant difference between OA and *Y*_*SR*_ for both CamCAN and Berlin. The optimal conduction delay significantly increases compared to OA in 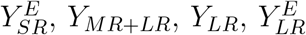 (CamCAN: *F* (77)=7.95, *p<*0.05 for 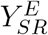; *F* (71)=12.63, *p<*0.05 for *Y*_*MR*+*LR*_; *F* (68)=5.73, *p<*0.05 for *Y*_*LR*_; *F* (74)=6.99, *p<*0.05 for 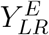 ; Berlin: *Z*=3.09, *p<*0.05 for 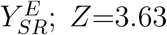, *p<*0.05 for *Z*; *Z*=2.88, *p<*0.05 for *Y*_*LR*_; *Z*=3.17, *p<*0.05 for 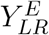) .

However, there is no significant difference between OA and *Y*_*SR*_ for both datasets, indicating that when the system has only SR connections, the conduction delay value is similar to that of the older cohorts. We find a consistent observation when comparing the coupling strength (and conduction delay) of older cohorts and the sub-graph solely connected via short-range fibers in both datasets.

## 4 Discussion

The study highlights the role of two sub-communities comprising short-range (SR) and long-range (LR) connections in aging-related compensatory mechanisms, particularly the role of the SR community in maintaining desired brain dynamics despite the decline in long-range fibers with age. We observe an overall loss of white matter tracts, especially in long-range fibers. However, despite this structural decline, the brain’s optimal working point and dynamical complexity remain unaltered as measured by metastability and entropy. This suggests an age-related adaptive mechanism of compensation. The large-scale brain network dynamics through simulations show that optimal global coupling strength and delay work together in a complementary fashion to maintain the brain’s optimal dynamics and preserve dynamic complexity with aging (indicated by metastability and entropy). We also observe that SR connections help enhance global coupling strength to compensate for the loss of long-range connections dynamically. SR and LR exceptions may not directly contribute to the coupling strength or conduction delay. Our observations on compensatory mechanisms driven separately by the short-range and long-range networks could help develop interventions to maintain optimal brain function in neurodevelopmental and aging-related neurodegenerative disorders.

### 4.1 Anatomical alterations with age

Age-associated loss of the brain’s anatomical connectivity has been of intense interest in many recent studies [Betzel et al., 2014, Lim et al., 2015, Perry et al., 2015]. Nevertheless, many of these studies have overlooked the impact on the short-range and long-range tract lengths, exceptions and their spatial distribution with age, which we quantified here and has been one of the key contributions of this work. To this end, we categorized three sub-community networks based on the white matter tract lengths, which include short-range (SR), mid-range (MR), and long-range (LR) connections. SR connections are the most common type of connection, accounting for approximately 80% of the total connection strength in the structural topology. MR and LR strengths accounted for 14% and 6% of the total fiber strength, respectively. White matter tract loss is estimated to be approximately 35% for the CamCAN dataset and 11% for the Berlin dataset in elderly individuals. On average, the total counts of SR tracts are reduced by approximately 31% in elderly subjects, while the total strength of MR and LR connections is reduced by 64% and 78%, respectively. These findings are consistent with recent works suggesting white matter connectivity loss is especially pronounced for the inter-hemispheric links [Puxeddu et al., 2020], causing the average length of the remaining tracts to decrease with age because the inter-hemispheric links are, on average, longer. Furthermore, we observe that the total tract length is reduced by 13% in the CamCAN dataset (2% in the Berlin dataset) in the elder group. Interestingly, it is seen that the total SR tract length (6%) is reduced more than MR and LR length (2% and 3%, respectively). Our results indicate that aging is associated with a significant loss of white matter connections, particularly in MR and LR connections, with a more noticeable decline in long-range connections. LR loss may be higher since they are highly metabolically demanding [Bullmore and Sporns, 2012] and require more energy to maintain, making them more vulnerable to age-related atrophy [Meijer et al., 2020].

### 4.2 Preserved dynamical complexity in the aging brain

The term ‘metastability’ was popularized by Scott Kelso, who drew inspiration from the works of Rodalfo Linas and Fransisco Varela to carry out fundamental brain-behavior experiments [Llinás, 1988, Kelso, 2012]. Their work shows metastability in the brain as a transient state where the whole-brain neural circuit can choose any attractor in response to external stimulus [Kelso, 2012]. Deco et al. further demonstrated that the resting brain operates at maximum metastability [Deco et al., 2017]. This state can be interpreted as a state in which the brain’s information processing capabilities are optimized, resulting in increased flexibility [Naik et al., 2017]. The dynamical working point of healthy aging has been suggested to lie within the regime of maximum metastability [Naik et al., 2017, Naskar et al., 2021]. Alterations in metastability have been observed in neuronal disorder, which is related to changes in dynamic repertories [Hellyer et al., 2015, Váša et al., 2015, Córdova-Palomera et al., 2017, Lee et al., 2018, Hancock et al., 2023], e.g., reduction in neural metastability linked to damage in the connectome and had behavioral implications [Hellyer et al., 2015], focal lesion could disrupt its anatomical architecture leading to alterations in metastability [Váša et al., 2015], decreased metastability in Alzheimer’s [Córdova-Palomera et al., 2017] and increased salience network metastability in schizophrenia [Lee et al., 2018,Hancock et al., 2023]. The dynamical measure of metastability can better indicate the overall dynamical complexity of the healthy aging brain.

Apart from metastability, we also quantify global entropy to quantify brain complexity with age. Entropy is a gauge of the nervous system’s ability to process information [Bergström and Nevanlinna, 1972, Fagerholm et al., 2023, Hancock et al., 2022]. It is a reliable metric for assessing brain function, reflecting its complexity and unpredictability. Higher neural entropy signifies complex and varied brain activity patterns, indicating improved adaptability in processing a wide range of information and making effective decisions in complicated tasks [Keshmiri, 2020]. On the other hand, lower neural entropy indicates more ordered or repetitive neural activities, which may reduce the brain’s ability to process diverse information but could enhance efficiency for specific, repetitive tasks. An optimized level of neural entropy, balancing entropy and redundancy in neural activity, might represent the ideal brain information processing state. The stability of entropy across two age groups suggests that the brain maintains its capacity to handle complex and varied information, adaptively responding to various cognitive demands.

The unchanged metastability and entropy with age highlight that the brain’s optimal dynamics remain stable, reflecting the resilience of its neural networks. It suggests that while aging may induce certain alterations in brain structure, an inherent mechanism persists to support cognitive flexibility and efficient information processing.

### 4.3 Complementary role of optimal coupling strength and delay

Notably, when the mean optimal coupling strength is low, the mean optimal conduction delay tends to increase, and vice versa. This pattern holds not only for young adults but also for older adults. In older adults, an increase in coupling strength may correspond to a decrease in conduction delay, indicating a compensatory mechanism that helps maintain optimal brain function. This finding is consistent across various subgraphs, including those connected via mid-range and long-range fibers. When the coupling strength is high, the conduction delay is low, suggesting that the brain adjusts these parameters to preserve its optimal working state. This complementary relationship between coupling strength and conduction delay appears to be a fundamental aspect of how the brain adapts to different structural and functional demands. Understanding this interplay is essential for elucidating the mechanisms of brain function and compensation in healthy aging and the context of neurodegenerative disorders.

### 4.4 Age-related compensatory mechanism

One crucial facet of brain plasticity lies in its ability to establish new connections [Murphy and Corbett, 2009] and adjust underlying biophysical parameters, such as neurotransmitter modulation [Saha et al., 2023], in response to damage in brain topology. This process is often called the brain’s neuro compensatory mechanism [Lövdén et al., 2010, Pathak et al., 2022, Petkoski et al., 2023, Saha et al., 2023]. The scaffolding process is present across the lifespan, as suggested in previous work, and involves using and developing complementary, alternative neural circuits to achieve a particular cognitive goal [Park and Reuter-Lorenz, 2009]. Hence, the behavioral performance in older adults is often hypothesized to depend on keeping some quantity invariant through compensation in another. The preservation of neuronal synchrony in aging through enhancing inter-areal coupling has been suggested previously by Pathak et al. (2022b). Here, we do not investigate a functional feature that is kept invariant. Instead, through a battery of dynamic features (metastability and entropy), we report that functional reorganization is associated with healthy aging. We demonstrate that the increase of global coupling combined with demyelination (change in conduction velocity) is necessary for this to occur in the brain model. On the other hand, the structural connectivity significantly decreases with aging, implying that the increased global coupling compensates for the lost structural connectivity consistent with recent findings [Pathak et al., 2022, Petkoski et al., 2023]. This is entirely plausible from the dynamical systems viewpoint, but it is unclear which biophysical mechanism could be operative for the shift in the global coupling strength. This is a purely phenomenological parameter, which has also been shown to be important in setting the working point in the case of epilepsy [Courtiol et al., 2020]. Despite this, based on personalized large-scale generative models in two independent datasets, our observations point towards a dynamical compensatory mechanism associated with aging, consistent with earlier reports by [Park and Reuter-Lorenz, 2009]. Despite significant structural loss, this mechanism is characterized by preserving dynamic complexity, as indicated by the unaltered metastability and entropy. Our results reveal that the aging brain adjusts parameters to maintain dynamic complexity despite fiber deterioration. A complementary interplay of global coupling strength and conduction delay could be one way to calibrate parameters to achieve optimal working states. Again, global coupling strength, which scales the local interaction strength among brain regions in older individuals, is significantly modulated and primarily contributes to the dynamical compensation, in line with prior observations that show that the strongest decrease with aging is observed for the inter-hemispheric tract lengths in aging [Petkoski et al., 2023, Pathak et al., 2022].

### 4.5 Role of SR and LR sub-communities on compensation

Further, we have highlighted the specific roles of the two sub-communities connected via short- and long-range tracts in modifying global interaction strength and conduction delay. We also investigate the impact of SR and LR exceptions in the aging brain, which has rarely been examined. Short-range tracts (low-cost connections) can remarkably reduce energy and resource consumption when forming, utilizing, and maintaining brain connections [Raichle and Mintun, 2006]. Moreover, they play a pivotal role in developmental disorders [Ouyang et al., 2017] and act as the most efficient communication pathways across the whole brain [Mišić et al., 2015]. The presence of both hypo- and hyper-connectivity in short-range fibers has been identified during early childhood in individuals with Autism Spectrum Disorder (ASD) [Rudie and Dapretto, 2013] and schizophrenia [McGlashan and Hoffman, 2000].

In older adults and those with Alzheimer’s Disease (AD), the impairment of local connectivity contributes to lower cognitive efficiency and results in higher compensatory neuronal activity [Gao et al., 2014]. Our results show no significant shift in the optimal coupling strength when comparing the sub-graph comprising solely short-range (SR) fibers with older adults (OA). This suggests that short-range connections modulate coupling strength in the aging brain to sustain the desired working point. The importance of SR connections on brain metastable dynamics is confirmed by Arvin et al. [Arvin et al., 2022].

The findings are consistent when comparing older adult cohorts’ coupling strength and conduction delay with the subgraphs solely connected via short-range fibers across both datasets. Specifically, in both the CamCAN and Berlin datasets, the conduction delay values for older adults are comparable to those observed in subgraphs with only short-range connections. This indicates that the dynamics of these connections are critical in understanding age-related changes in brain function.

On the other hand, long-range white matter tracts, characterized by their myelinated fibers, play a crucial role in carrying long distant synapses across the brain that facilitate efficient and effective communication supporting various cognitive functions [Sporns and Zwi, 2004]. The scientific literature extensively investigates age-related loss of long-range white matter tracts and its subsequent impact on conduction velocity [Sullivan and Pfefferbaum, 2006,Bennett et al., 2010]. The importance of long-range exceptions for functional connectivity is also highlighted in previous studies [Deco et al., 2021]. The observed decrease in LR exceptions with age implies potential effects on functional connectivity and underlying dynamics. However, our findings show that the brain’s metastable dynamics, where it operates, are preserved. This preservation is likely due to the rescaling of coupling strength and conduction delay, compensating for losing LR exceptions. The enhancement of coupling strength in the subgraph connected via short-range connections may reflect this dynamic compensation. We provide a parsimonious explanation for the systematic enhancement of coupling strength with age by demonstrating a systematic reduction in long-range connectivity and the creation of new clusters with increased local short-range connectivity. This supports partial synchronization in the complex network. One possible reason for the lack of difference in brain dynamics compared to young adults could be the accelerated degeneration of long-range neural fibers, resulting in over-activation at the whole brain level.

### 4.6 Limitations

Several limitations in our study need to be addressed in future research. To begin with, the Kumamoto model captures the emerged phase dynamics of the neural signals in the whole brain. Instead, more realistic models could be utilized to gain further detailed insights into the underlying alterations in excitation-inhibition balance and neurotransmitter levels [Vattikonda et al., 2016, Naskar et al., 2021]. Furthermore, the network investigation was based on a coarse parcellation scheme comprising only 68 brain regions. However, our results will likely be validated using other high-resolution parcellations. Moreover, there are inherent limitations in tractography measures using diffusion MRI techniques, such as the failure to identify existing tracts in cross-hemispheric connectivity. It could be challenging to resolve accurately as the uncertainty in a streamlined location increases with the length of the fiber tracts. Additionally, there are systematic differences in brain size, grey matter atrophy, and white matter hyperintensities with aging. However, little is known from existing literature on incorporating them as free parameters in the model for all the network simulations. Future studies can be performed to address this issue. Finally, it should be noted that while we use the Kuramoto order parameter, quantifying metastability captures the whole brain’s dynamical repertoire, which only measures inphase cohesion and its deviations. Other cohesion patterns, such as anti-phase or anti-synchrony, and their fluctuations are only partially captured here.

### 4.7 Conclusion

This study sheds light on the mechanisms underlying dynamical compensation in aging brains. We provide a detailed explanation of the critical role of short-range connections in the face of declining long-range connections. Our key findings are twofold: 1) Optimal coupling strength and delay act in concert to maintain the brain’s stable yet adaptable dynamics across ages and brain regions. 2) Short-range connections adjust their strength, preserving the brain’s optimal working (metastable) state despite aging. This adaptation ensures functional integrity even with declining long-range connections. While traditionally seen as essential for communication, decreasing long-range connections necessitates a compensatory role for short-range ones. Integrating detailed subgraph analysis, our novel methodology offers a unique framework to investigate the intricate relationship between brain structure and its complex dynamics. This approach holds significant promise for understanding compensatory mechanisms in neurological disorders and potentially informing the development of future therapeutic strategies. By paving the way for exploring brain function in healthy aging and pathological conditions, this research lays a groundbreaking foundation for future advancements in neuroscience.

## Acknowledgments

We acknowledge the generous support of NBRC Core funds and the Computing facility. For simulations, resources from Neuroscience Gateway [Sivagnanam et al., 2013] were used. SS is supported by NPDF, SERB-DST, India, Award ID: PDF/2021/000585. AB acknowledges NBRC Flagship program, DBT, India, Award ID: BT/MED-III/NBRC/Flagship/Flagship2019. AB is supported by the Ministry of Youth Affairs and Sports, India, Award ID: F.NO.K-15015/42/2018/SP-V. DR is funded by Ramalingaswami Fellowship, DBT, India, Award ID: BT/RLF/Re-entry/07/2014 and DST, India, Award ID: SR/CSRI/21/2016.

## Author contribution

PC: Data accusation; Conceptualization; Investigation; Methodology; Visualization; Writing original draft, Editing. SS: Conceptualization; Investigation; Methodology; Visualization; Writing original draft; Editing. GD: Methodology; Editing. AB: Methodology; Visualization; Funding acquisition; Supervision; Project Administration; Resources; Editing. DR: Conceptualization; Methodology; Visualization; Funding acquisition; Supervision; Project Administration; Resources; Writing original draft; Editing.

